# Structural basis for terminal loop recognition and processing of pri-miRNA-18a by hnRNP A1

**DOI:** 10.1101/178855

**Authors:** Hamed Kooshapur, Nila Roy Choudhury, Bernd Simon, Max Mühlbauer, Alexander Jussupow, Noemi Fernandez, Andre Dallmann, Frank Gabel, Carlo Camilloni, Gracjan Michlewski, Javier F. Caceres, Michael Sattler

**Author notes:** Correspondence and requests for materials should be addressed to G.M; J.F.C. or M.S.

## Abstract

Post-transcriptional mechanisms play a predominant role in the control of microRNA (miRNA) production. Recognition of the terminal loop of precursor miRNAs by RNA-binding proteins (RBPs) influences their processing; however, the mechanistic and structural basis for how levels of individual or subsets of miRNAs are regulated is mostly unexplored. We previously described a role for hnRNP A1, an RBP implicated in many aspects of RNA processing, as an auxiliary factor that promotes the Microprocessor-mediated processing of pri-mir-18a. Here, we reveal the mechanistic basis for this stimulatory role of hnRNP A1 by combining integrative structural biology with biochemical and functional assays. We demonstrate that hnRNP A1 forms a 1:1 complex with pri-mir-18a that involves binding of both RNA recognition motifs (RRMs) to cognate RNA sequence motifs in the conserved terminal loop of pri-mir-18a. Terminal loop binding induces an allosteric destabilization of base-pairing in the pri-mir-18a stem that promotes its down-stream processing. Our results highlight terminal loop RNA recognition by RNA-binding proteins as a general principle of miRNA biogenesis and regulation.

---

MicroRNAs (miRNAs, miRs) are a class of highly conserved small non-coding RNAs that play a crucial role in the regulation of gene expression. They are involved in a variety of biological processes including cell growth, proliferation and differentiation^1^. Mature miRNAs are generated by two RNA cleavage steps involving nuclear and cytoplasmic RNase III enzymes (Drosha and Dicer, respectively). Primary miRNA (pri-miRNA or pri-mir) transcripts are cropped by the Microprocessor complex (comprising Drosha and DGCR8) in the nucleus forming ~70 nucleotide (nt) stem-loop precursor miRNAs (pre-miRNAs or pre-mir), which, following export to the cytoplasm, are further processed by Dicer into mature miRNAs (reviewed in^2^). Many miRNA genes in higher organisms are transcribed together as a cluster^3^. A prototypical example is the miR-17-92 cluster that is encoded as an intronic polycistron on chromosome 13 in humans. This cluster encodes six individual miRNAs that are highly conserved in vertebrates (miR-17, miR-18a, miR-19a, miR-20a, miR-19b-1, miR-92a-1, reviewed in^4^). The miR-17-92 cluster is frequently amplified and overexpressed in human cancers; hence, it is also referred to as OncomiR-1. Its oncogenic role was confirmed in a mouse model of B cell lymphoma^5^. Furthermore, its targeted deletion is associated with developmental defects in mouse model systems^6^.

The biogenesis of miRNAs is tightly regulated and results in tissue- and developmental-specific expression patterns of miRNAs^7^. A number of specific RNA-binding proteins (RBPs) have recently emerged as important post-transcriptional regulators of miRNA processing. However, very little is known about their mechanism of action. Previously, we identified heterogenous nuclear ribonucleoprotein A1 (hnRNP A1) as a factor, which positively regulates the processing of miRNA-18a primary transcript (pri-mir-18a) by making specific contacts to the terminal loop of the RNA^8,9^ (Fig. 1a,b). HnRNP A1 is a highly abundant RBP that has been implicated in diverse cellular functions related to RNA processing, including alternative splicing regulation^10–12^, mRNA export^13,14^, IRES (internal ribosome entry site)-mediated translation^15,16^, mRNA stability^17,16^ and telomere maintenance^18,19^.

**Fig. 1.**
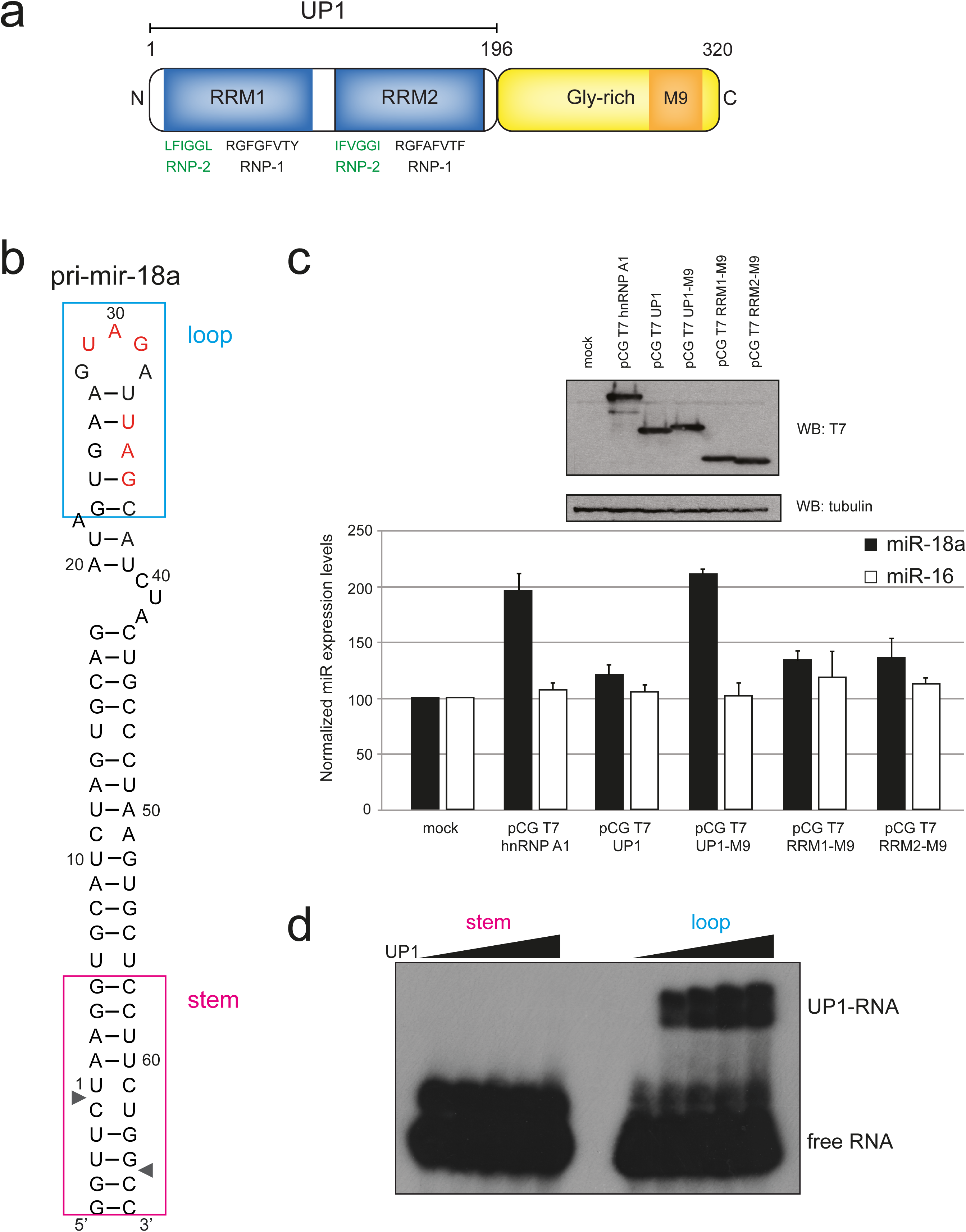
The tandem RRMs (UP1) of hnRNP A1 are necessary and sufficient to promote pri-mir-18a biogenesis in living cells. (**a**) Domain structure of human hnRNP A1. The sequences of conserved RNP-1 and RNP-2 motifs in the RRM domains are indicated. M9 is a transport signal linked with both nuclear import and export of this protein. (**b**) Secondary structure of pri-mir-18a RNA based on footprinting analysis^9^. Regions corresponding to the terminal loop and stem are boxed. The cleavage sites for Microprocessor (Drosha/DGCR8) are indicated by arrowheads. (**c**) Effect of transiently transfected epitope-tagged T7-hnRNP A1, UP1, UP1-M9, RRM1-M9 and RRM2-M9 in the processing of pri-mir-18a in HeLa cells in culture. Processing of pri-mir-16 was included as a control (white bars). The upper panel shows the level of expression of T7 epitope-tagged hnRNP A1 WT or constructs expressing individual domains (RRM1 or RRM2) or the UP1 fragment (tandem RRM1-RRM2). An M9 sequence was included to direct the nuclear localization of the UP1, RRM1 and RRM2 constructs. (**d**) Electrophoretic mobility shift assay (EMSA) of UP1 in complex with pri-mir-18a loop and stem RNAs.

Here, we have combined an integrative structural biology approach with biochemical and functional assays to provide mechanistic insights into the role of hnRNP A1 in stimulating primir-18a processing. We show that hnRNP A1 forms a 1:1 complex with pri-mir-18a in solution, with the recognition of two UAG motifs by the tandem RRM domains of hnRNP A1 revealed by a high-resolution crystal structure. NMR and biophysical data show that high-affinity binding involves recognition of two UAG motifs in the pri-mir-18a terminal loop and the proximal stem region. Notably, binding to the terminal loop induces an allosteric destabilization of base-pairing in the pri-mir-18a stem that promote its processing. These findings may serve as a paradigm for the regulation of miRNA processing by the recognition of the terminal loop by RBPs.

## Results

### Nuclear localized UP1 is necessary and sufficient for stimulating pri-mir-18a processing

We have previously shown that hnRNP A1 acts as an auxiliary factor for miRNA biogenesis, by binding to pri-mir-18a and inducing a relaxation at its lower stem creating a more favorable cleavage site for Drosha^9^. However, the underlying molecular mechanisms are unknown. HnRNP A1 has two RNA recognition motif (RRM) domains, each harboring conserved RNP-1 and RNP-2 submotifs, and a C-terminal flexible glycine-rich tail, which includes the M9 sequence, responsible for nuclear import and export^20,13.^ The RNA-binding region of hnRNP A1, comprising the tandem RRM1-RRM2 domains, is referred to as UP1 (Unwinding Protein 1)^21^ (Fig. 1a). To identify which regions in hnRNP A1 are required for stimulating pri-mir-18a processing in living cells we used an *in vivo* processing assay. For this, several N-terminal T7-tagged hnRNP A1 constructs were transiently overexpressed in HeLa cells and the level of mature miR-18a was analyzed by qRT-PCR. We found that overexpression of full-length hnRNP A1 results in a ~2-fold increase in the levels of mature miRNA-18a, whereas UP1, comprising both RRMs but lacking the M9 sequence, has no effect – most likely due to its cytoplasmic localization (Fig. 1c; Supplementary Fig. 1a, b; Supplementary Table 1). This was confirmed by transient expression of UP1-M9, where UP1 is fused to the M9 sequence that directs nuclear localization of hnRNPA1^20^. UP1-M9 localized exclusively to the nucleus (Supplementary Fig. 1b; Supplementary Table 1) and, importantly, displayed similar activity as full-length hnRNP A1 in stimulating miR-18a production *in vivo* (Fig. 1c). By contrast, RRM1-M9 and RRM2-M9 that partially localize to the nucleus (Supplementary Fig. 1b; Supplementary Table 1) did not increase miR-18a levels, showing that nuclear localized UP1, i.e. comprising both RRMs of hnRNP A1, is required for function (Fig. 1c).

**Table 1.** Isothermal titration calorimetry of protein-RNA interactions

| Binding partners | N<br>(protein:RNA) | $K_D$<br>(M) | $\Delta H$<br>(cal/mol) | $\Delta S$<br>(cal/mol/deg) |
| --- | --- | --- | --- | --- |
| RRM1 + 7-mer<br>(AGUAGAU) | 1.01 ± 0.02 | (20.4 ± 1.06)<br>x 10<sup>-6</sup> | -1.988 x 10<sup>4</sup> ±<br>646.1 | -45.2 |
| RRM2 + 7-mer<br>(AGUAGAU) | 1.09 ± 0.00 | (6.8 ± 0.14)<br>x 10<sup>-6</sup> | -1.645 x 10<sup>4</sup> ±<br>62.21 | -31.6 |
| UP1 + 7-mer (AGUAGAU) | 0.75 ± 0.00 | (3.4 ± 0.12)<br>x 10<sup>-6</sup> | -3.530 x 10<sup>4</sup> ±<br>353.6 | -93.4 |
| UP1 + 17-mer(A35C) | 1.07 ± 0.01 | (3.1 ± 0.19)<br>x 10<sup>-6</sup> | -1.979 x 10<sup>4</sup> ±<br>314.7 | -41.2 |
| UP1 + pri-mir-18a | 1.35 ± 0.00 | (147.7 ± 8.9)<br>x 10<sup>-9</sup> | -3.522 x 10<sup>4</sup> ±<br>151.2 | -86.8 |
| UP1 + 12-mer<br>(AGUAGAUUAGCA) | 1.01 ± 0.00 | (15.5 ± 3.4)<br>x 10<sup>-9</sup> | -3.809 x 10<sup>4</sup> ±<br>200.4 | -92.0 |
| UP1 + 12-mer-mut1<br>(AGUuuAUUAGCA) | 1.06 ± 0.00 | (154.1 ± 11)<br>x 10<sup>-9</sup> | -2.974 x 10<sup>4</sup> ±<br>164.2 | -68.6 |
| UP1 + 12-mer-mut2<br>(AGUAGAUUuuCA) | 1.08 ± 0.00 | (330.0 ±<br>26.47) x 10<sup>-9</sup> | -2.449 x 10<sup>4</sup> ±<br>208.2 | -52.5 |
| UP1 + 12-mer-mut3<br>(uuUAGAUUAGCA) | 0.98 ± 0.00 | (8.33 ± 1.35)<br>x 10<sup>-9</sup> | -4.102 x 10<sup>4</sup> ±<br>197.9 | -101 |
| UP1 + 10-mer<br>(UAGAUUAGCA) | 1.07 ± 0.00 | (19.12 ±<br>6.73) x 10<sup>-9</sup> | -3.938 x 10<sup>4</sup> ±<br>396.4 | -96.8 |
| UP1(R75E/R88A) + 12-mer<br>(AGUAGAUUAGCA) | 1.08 ± 0.00 | (19.01 ±<br>2.05) x 10<sup>-9</sup> | -3.841 x 10<sup>4</sup> ±<br>158.3 | -93.5 |
| UP1(R75E/R88E) + 12-mer<br>(AGUAGAUUAGCA) | 1.1 ± 0.00 | (38.8 ± 3.44)<br>x 10<sup>-9</sup> | -3.68 x 10<sup>4</sup> ±<br>160.0 | -89.5 |

### UP1 specifically recognizes the loop region of pri-mir-18a

We next wanted to determine the regions of pri-mir-18a that are recognized by hnRNP A1. To this end, we performed electro-mobility shift assays (EMSA) with RNA variants corresponding to the terminal loop and the stem of pri-mir-18a with UP1 (Fig. 1b), which has been shown to recapitulate most of the functions of full-length hnRNP A1 *in vitro*^21^ (Fig. 1d). We observed that UP1 specifically binds to the terminal loop RNA, whereas no binding to the stem RNA was detected even at higher protein-to-RNA ratios. Single RRM1 and RRM2 domains do not show any detectable RNA-binding activity in this assay (Supplementary Fig. 1c). Altogether, these data show that i) UP1 binds specifically to the terminal loop region of pri-mir-18a, and ii) both RRM domains of UP1 are required for high affinity RNA binding, indicating that they bind cooperatively.

### Identification of a minimal UP1-binding sequence in the pri-mir-18a terminal loop

To provide a quantitative analysis of binding affinities of UP1 with pri-mir-18a, we performed isothermal titration calorimetry (ITC) experiments. Pri-mir-18a 71-mer RNA binds to UP1 with a dissociation constant (*K*_D_) of 147 nM (Fig. 2a). To identify a minimal RNA region required for efficient UP1 binding, we tested RNA fragments of various sizes for binding to UP1 and individual RRMs by ITC (Table 1; Fig. 2a; Supplementary Fig. 2a). A 7-mer oligonucleotide (5’ AG<u>UAG</u>AU 3’) corresponding to the terminal loop of pri-mir-18a harbors an UAG motif, which is known to be recognized by hnRNP A1^22^. This 7-mer single-stranded RNA binds to the individual RRM1 and RRM2 domains with low micromolar affinity (*K*_D_ = 20.4 µM and 6.9 µM, respectively) forming 1:1 complexes (Table 1; Supplementary Fig. 2a). Binding of the 7-mer to UP1 has a *K*_D_ of 3.4 µM (Table 1; Fig. 2a, left panel), suggesting that each RRM domain in UP1 can recognize the 7-mer RNA. Notably, a single-stranded 12-mer oligonucleotide (5’ AG<u>UAG</u>AU<u>UAG</u>CA 3’) derived from the pri-mir-18a terminal loop and flanking sequences (Fig. 1b) shows more than 200-fold higher affinity (*K*_D_ = 15.5 nM with a 1:1 stoichiometry) to UP1 (Fig. 2a, middle panel). The 12-mer RNA harbors two UAG motifs suggesting that each of these can be recognized by one of the RRMs in a cooperative manner. This is evident from the very large increase in binding affinity compared to the binding of the 7-mer to UP1. It is remarkable that binding of UP1 to this single-stranded 12-mer RNA is at least 10-fold stronger than binding to full-length pri-mir-18a (*K*_D_ = 15.5 nM vs. 147 nM, respectively) (Fig. 2a, middle and right panel, respectively). This may be related to the observation that in the pri-mir-18a stem-loop the second UAG motif is predicted to be base-paired in the loop-proximal stem and thus not freely accessible (Fig. 1b). Binding of UP1 to pri-mir-18a may thus require opening (melting) of these base-pairs, whereas in the single-stranded 12-mer RNA both UAG binding sites are readily available for interaction with UP1 (see below), thus the higher affinity. We conclude that the 12-mer RNA is a high-affinity UP1-binding sequence where both RRM domains of UP1 recognize UAG motifs. The RNA recognition features in the 12-mer are expected to represent the interaction of UP1 with the pri-mir-18a. A summary of thermodynamic parameters of the hnRNPA1-RNA interactions is given in Table 1.

**Fig. 2.**
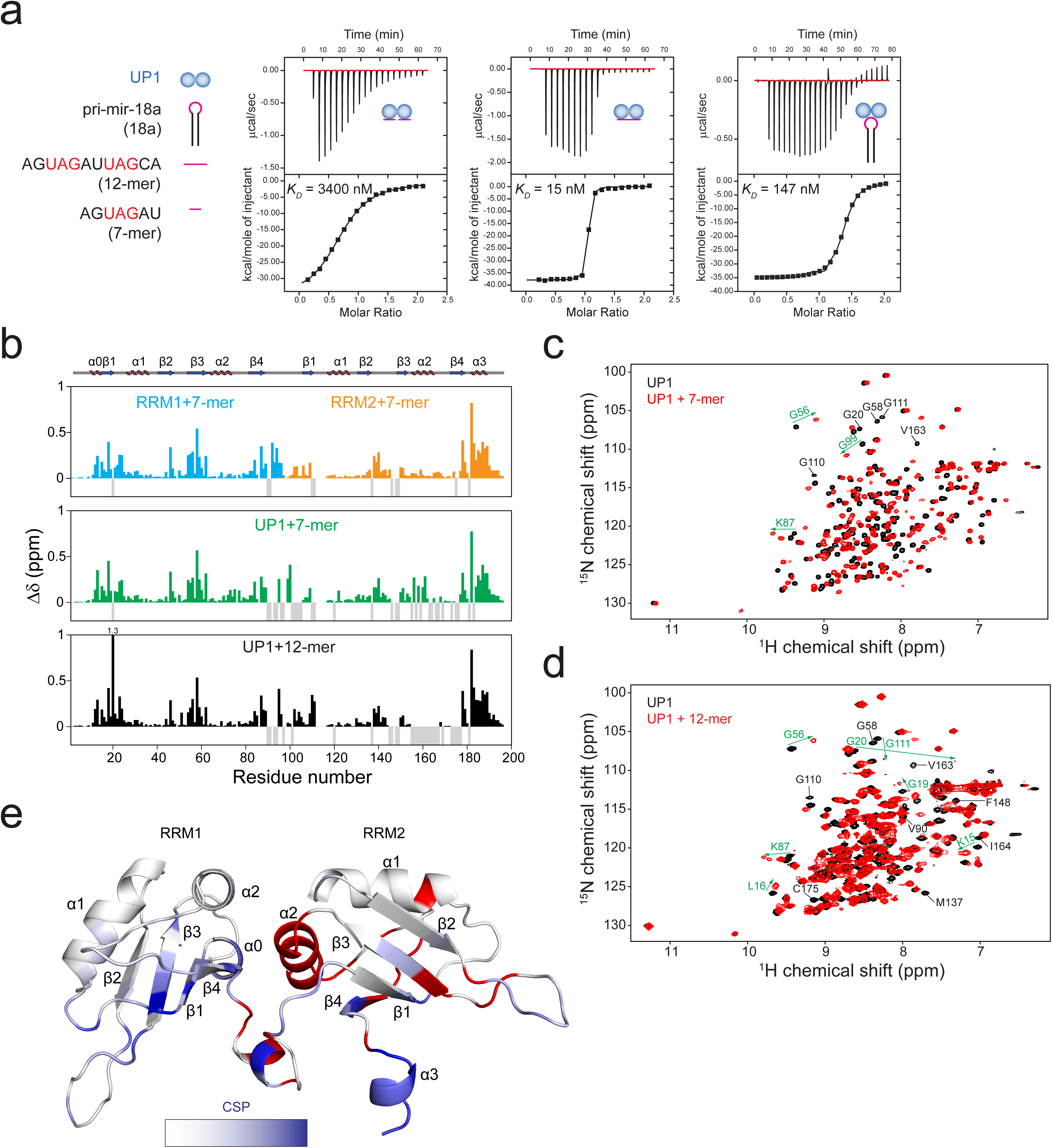
Biophysical characterization of UP1-RNA interactions. (**a**) ITC of the binding of UP1 to 7-mer, 12-mer and pri-mir-18a RNAs. *KD* values are indicated. The sequences of 7-mer and 12-mer oli.gonucleotides derived from the terminal loop of pri-mir-18a are shown on the left. (b) Combined ^1^H and ^15^N chemical shift perturbations (CSPs) of RRM1/7-mer, RRM2/7-mer, UP1/7-mer and UP1/12-mer are plotted against the residue number. Secondary structure elements are shown above the plot. The gaps in the graph are proline residues; negative grey bars represent residues that could not be assigned due to line-broadening. (**c, d**) ^1^H, ^15^N correlation spectra of UP1, free (black) and in the presence of either the (**c**) 7-mer or (**d**) the 12-mer RNA (red). Selected residues experiencing large chemical shift perturbations and line-broadening upon RNA binding are labeled in green and black, respectively. (**e**) The CSPs are mapped onto the structure of UP1 (gradient of white to blue indicates weak to strong CSPs). Residues corresponding to amide signals that are exchange-broadened in the RNA-bound spectra are colored red.

### NMR analysis of UP1-RNA interactions

We next characterized the binding interface of UP1 with various RNA ligands derived from pri-mir-18a using NMR titration experiments. Addition of the 7-mer RNA harboring one UAG motif causes extensive chemical shift perturbations (CSPs) in RRM1 and RRM2 constructs, thus demonstrating that each of the RRMs can interact with the 7-mer RNA (Fig. 2b). The CSP pattern obtained upon titration of the tandem RRM domains in UP1 with the 7-mer RNA is very similar to the one obtained for individual RRM domains (Fig. 2b, c). This shows that the recognition of the RNA in the isolated RRM domains and in the context of the UP1 construct is very similar. Similarly, the 12-mer RNA harboring two UAG motifs induces large CSPs in both RRM domains of UP1 (Fig. 2d) that are comparable to those obtained at saturating levels of the 7-mer. The CSPs map to the canonical RNA binding surface on the β-sheets of the two RRM domains (Fig. 2e). In addition, strong CSPs are observed for residues in the C-terminal region of RRM2. This region is flexible in free UP1 but upon RNA binding forms an additional helix (α3), which is not present in the free protein^23^ (Supplementary Fig. 3a, b), suggesting that helix α3 is induced and stabilized upon RNA binding. Interestingly, a number of NMR signals that correspond to residues in RRM2 and the RRM1-RRM2 linker are severely broadened in the RNA-bound spectrum, suggesting dynamics on the µs-ms time-scale. The affected residues map to the interface between the RRM1 and RRM2 domains (Fig. 2e), suggesting that some conformational dynamics and adaptation of this domain interface is associated with RNA binding.

**Fig. 3.**
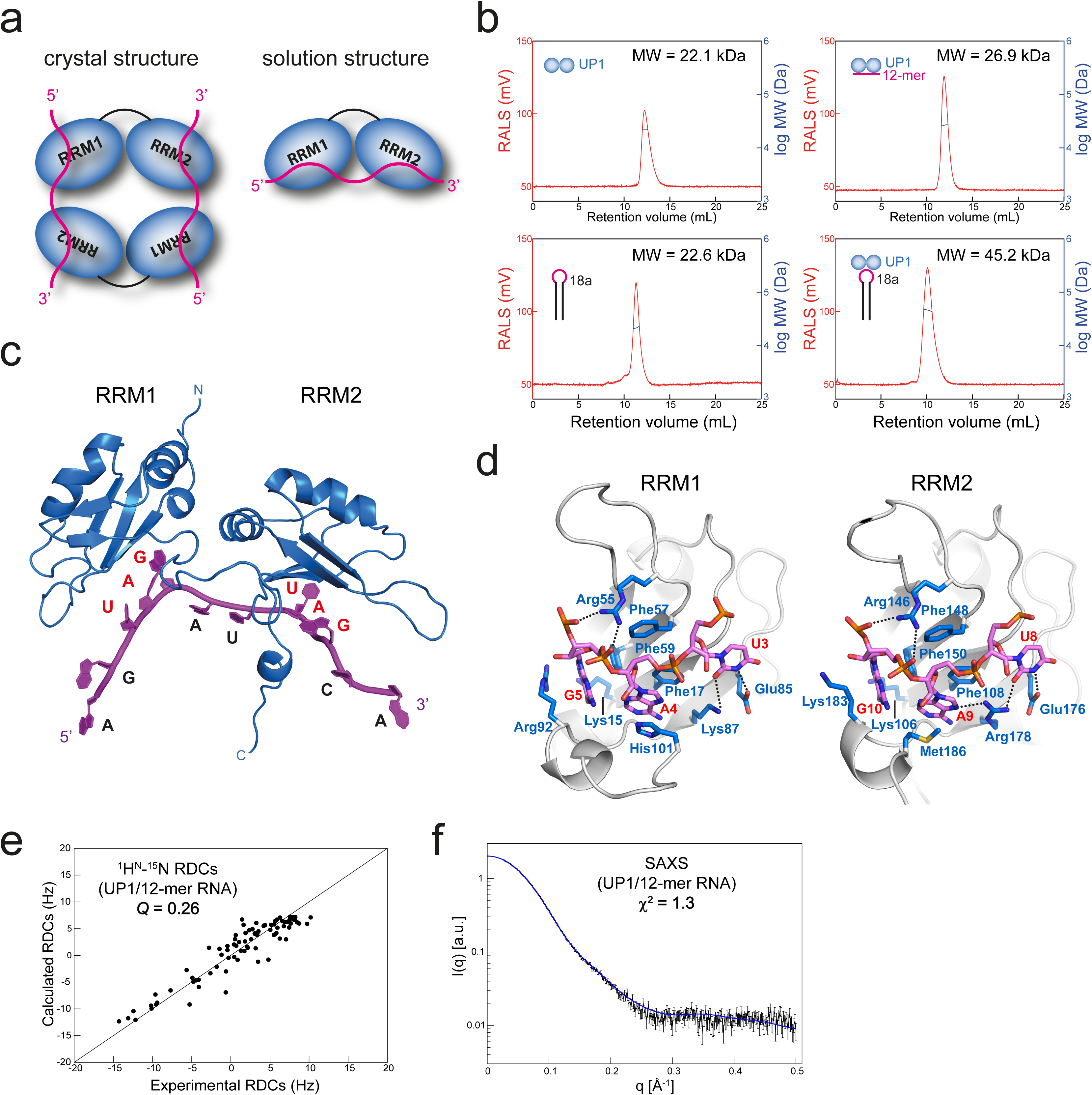
Structure of the UP1/12-mer complex. (**a**) Schematic representation of the 2:2 UP1/12-mer RNA complex observed in the crystal structure (Supplementary Fig. 4a) and the proposed 1:1 complex in solution. (**b**) Static light scattering (SLS) profiles of UP1, pri-mir-18a, UP1/12-mer and UP1/pri-mir-18a. The molecular weight (MW) obtained from SLS is indicated. UP1 forms a 1:1 complex with both pri-mir-18a and the 12-mer oligonucleotide. (**c**) Structural model of the 1:1 UP1/12-mer complex, where each RRM domain recognizes a UAG motif in the 12-mer RNA (magenta) forming a 1:1 complex. (**d**) Structural details of recognition of the UAG motif by RRM1 and RRM2 domains. Side-chain of selected residues involved in the interaction are indicated. (**e, f**) Correlation of experimental ^1^H-^15^N residual dipolar couplings (RDCs) (**e**) and small angle X-ray scattering (SAXS) data (**f**) *vs.* those calculated from the structural model shown in (c).

### Structural basis for the recognition of the pri-mir-18a terminal loop by UP1

To gain insight into the molecular details of pri-mir-18a recognition by UP1, we determined the crystal structure of the UP1/12-mer RNA complex at 2.5 Å resolution (Table 2; Fig. 3a; Supplementary Fig. 4a). Surprisingly, the crystal structure of the UP1/12-mer RNA complex exhibits two molecules of UP1 and two RNA chains in the asymmetric unit in a 2:2 stoichiometry (Supplementary Fig. 4a), similar to a previously reported structure of UP1 with single-stranded telomeric DNA^24^. As this peculiar 2:2 stoichiometry most likely does not represent the UP1:RNA complex in solution, we analyzed UP1, RNA and the protein-RNA complexes by static light scattering (SLS) (Fig. 3b). Both UP1 and pri-mir-18a alone behave as single species with a molecular weight corresponding to respective monomeric conformations. Importantly, the molecular weight obtained for the UP1/12-mer complex (22.4 kDa) indicates a 1:1 complex and demonstrates that the 2:2 stoichiometry observed in the crystal structure is an artifact induced by the crystal environment. Notably, the molecular weight obtained for the UP1/pri-mir-18a complex (45 kDa) is fully consistent with the formation of a 1:1 complex (Fig. 3a, b).

**Table 2.** Crystallographic data collection and refinement statistics

|  | UP1/12-mer |
| --- | --- |
| **Data collection** |  |
| Space group | P6222 |
| Cell dimensions |  |
| $a, b, c$ (Å) | 127.03, 127.03, 147.06 |
| $\alpha, \beta, \gamma$ (°) | 90, 90, 120 |
| Resolution (Å) | 48-2.5 |
|  | (2.6-2.5) |
| $R_{\text{merge}}$ | 0.09 (0.98) |
| $I / \sigma I$ | 20.54 (2.09) |
| Completeness (%) | 99.9 (99.1) |
| Redundancy | 9.9 (9.1) |
| **Refinement** |  |
| Resolution (Å) | 48-2.5 |
| No. reflections | 24,897 |
| $R_{\text{work}} / R_{\text{free}}$ | 0.19 / 0.23 |
| No. atoms | 3411 |
| Protein | 2880 |
| RNA | 455 |
| Water | 72 |
| $B$-factors | |
| Protein | 72.81 |
| RNA | 87.93 |
| Water | 48.54 |
| R.m.s. deviations |  |
| Bond lengths (Å) | 0.009 |
| Bond angles (°) | 1.25 |

The dataset is from a single crystal. Values in parentheses are for highest-resolution shell.

**Fig. 4.**
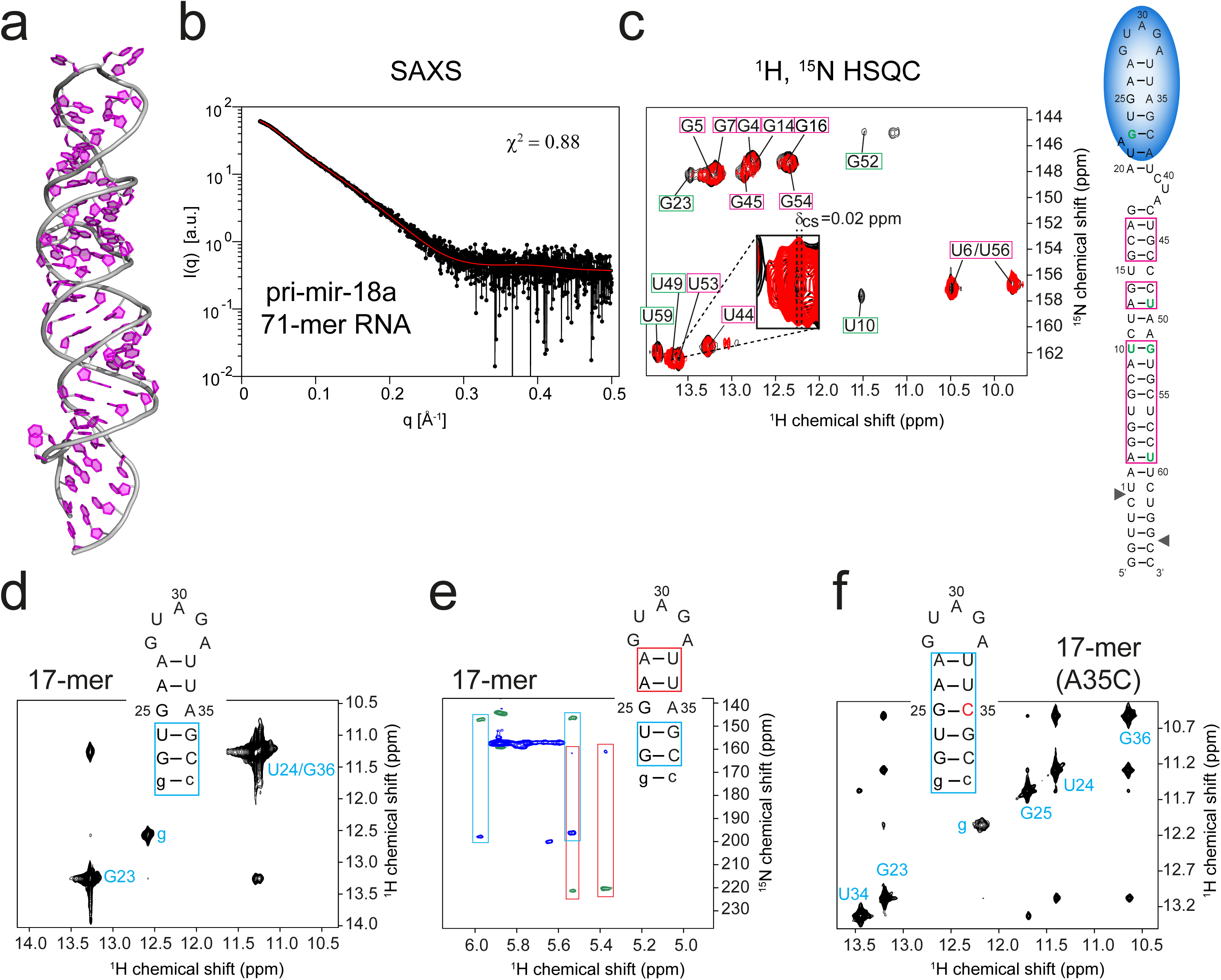
Structural analysis of pri-mir-18a 71-mer RNA. (**a**) A structural model of pri-mir-18a 71-mer RNA obtained using the MC-Fold/MC-Sym server. (**b**) Experimental and predicted SAXS data for the pri-mir-18a model are shown in black and red, respectively. (**c**) NMR analysis of the 71-mer pri-mir-18a RNA. ^1^H, ^15^N-HSQC of ^13^C, ^15^N-labeled pri-mir-18a in the absence (black) and presence (red) of UP1. Imino signals observed indicating stable base-pairing are shown on the secondary structure of pri-mir-18a (magenta boxes). Nucleotides undergoing large perturbations upon UP1 binding are highlighted with green boxes and green letters on the right. (**d**) 2D-imino NOESY and (**e**) H5 correlated HNN experiment of the 17-mer RNA derived from the terminal loop and flanking stem of pri-mir-18a. Base pairs confirmed by these experiments are boxed in blue and red on the secondary structure of the 17-mer RNA. The first G:C base pair, which is not part of the native sequence, is shown in lower case. (**f**) 2D-imino NOESY of the 17-mer(A35C) RNA. The mutated nucleotide is shown in red. Base pairs confirmed by the detection of the imino correlation are boxed.

The crystal structure reveals that each RRM domain in UP1 specifically recognizes one UAG motif in the RNA. Although the stoichiometry does not reflect the solution conformation, the RNA contacts are expected to be conserved, consistent with the NMR titrations. Each UAG motif is recognized by contacts mainly through conserved RNP motif residues on the β-sheets (Fig. 3c, d), which resembles the recognition of TAG in the UP1-telomeric DNA complex^24^ and recently reported structures with RNA^25,26.^ Two conserved aromatic residues in RRM1, Phe17 (RNP-2 motif residue located on β1) and Phe59 (RNP-1 motif residue located on β3), are involved in stacking interactions with the bases of A4 and G5, respectively (Fig. 3d). A third aromatic residue, Phe57 (RNP-1 motif residue located on β3), interacts with the ribose rings of A4 and G5. Similarly, in RRM2 Phe108 (RNP-2 motif residue located on β1) and Phe150 (RNP-1 motif residue located on β3) stack with the bases of A9 and G10, respectively, whereas Phe148 (RNP-1 motif residue located on β3) makes contacts with the sugar rings of A9 and G10. In addition to the stacking interactions with RNP residues (*vide infra*), the central adenosine in the UAG motifs is specifically recognized by hydrogen bonds of its exocyclic NH2 group with the main chain carbonyl oxygen of residues Arg88 and Lys179 in RRM1 and RRM2, respectively. A positively charged residue in each domain, Arg55 in RRM1 and Arg146 in RRM2, makes electrostatic interactions with the phosphate backbone of the AG dinucleotide. Two charged residues in each domain, Glu85 and Lys87 in RRM1 and Glu176 and Arg178 in RRM2, make specific contacts to the uridines in the UAG motifs (U3 and U8), while another charged residue, Lys15 in RRM1 and Lys106 in RRM2, interacts with G5 and G10, respectively (Fig. 3d), thereby specifying the U and G residues in the UAG motif. The mode of RNA recognition by RRM1 and RRM2 is very similar; in each domain an AG dinucleotide is sandwiched between the β-sheet surface and a C-terminal helix (Fig. 3d).

The two RRM domains of hnRNP A1 are connected by an approximately 17-residue linker, which is evolutionary conserved both in terms of sequence and length^27^ (Supplementary Fig. 3e). Residues in this linker are highly flexible as determined by NMR relaxation data (Supplementary Fig. 3a). The two RRM domains interact with each other and this intramolecular domain interface is stabilized by two conserved salt-bridges (Arg75-Asp155 and Arg88-Asp157) as well as a small cluster of hydrophobic residues in the interface. Virtually the same domain arrangement and salt bridges are observed in the monomeric solution structure of UP1 free^23^ and a crystal structure bound to a short RNA^25^ (Supplementary Fig. 4d).

To confirm the domain arrangement of RNA-bound UP1 in solution, we measured NMR paramagnetic relaxation enhancements (PRE) for UP1 spin-labeled at position 66 (UP1 Glu66Cys mutant). The PRE data provide long-range (up to ~20 Å) distance information and can thus report on domain/domain arrangements^28–30^ (Supplementary Fig. 4b). The PRE profiles of free and RNA-bound form of UP1 are similar, suggesting that the domain arrangement does not change significantly in the presence of the 12-mer RNA.

To obtain additional restraints to determine a structural model of the 1:1 UP1/12-mer RNA complex in solution we measured residual dipolar couplings (RDCs) and small angle X-ray scattering (SAXS) on the UP1/12-mer RNA complex (Fig. 3e, f; Supplementary Table 2). A single protein monomer from our crystal structure was used as template and a single RNA molecule was initially posed constraining the recognition of the two UAG motifs in the 12-mer RNA (Fig. 3a). SAXS and RDC data were then used as restraints in a molecular dynamics simulation together with the distances of the Arg75-Asp155 and Arg88-Asp157 salt-bridges as observed in the crystal structure. The refinement shows that a 1:1 complex is fully compatible with the data, where the arrangement of RRM1 and RRM2 is similar to the corresponding interface in the crystal structure (coordinate rmsd of 2.8 Å calculated over all the backbone atoms excluding linker residues) with only minor conformational changes in the domain/domain interface (Fig. 3d; Supplementary Fig. 4a-d). The structural model is in excellent agreement with the NMR and SAXS data (Fig. 3e, f).

### Validation of the UP1/12-mer RNA structural model

The structural model of the 1:1 UP1/12-mer RNA complex was confirmed by mutational analysis of protein and RNA. ITC data with 12-mer RNAs where the first or second UAG motif has been replaced by UUU, 12-mer-mut1: AG<u>UUU</u>AUUAGCA and 12-mer-mut2: AGUAGAU<u>UUU</u>CA show 10-fold and 20-fold (*K*_D_ = 154 nM and 330 nM) reduced binding affinity, respectively, compared to the wildtype sequence (Table 1; Supplementary Fig. 2b). This demonstrates that both motifs are recognized by the protein. Additional RNA variants with an AG→UU mutation or lacking the initial AG dinucleotide have the same binding affinity as the wildtype 12-mer RNA (Table 1; Supplementary Fig. 2b). This shows that UP1 has a preference for the recognition of two neighboring UAG motifs.

Further, the domain interface in UP1 was probed by mutations that are expected to disrupt the two salt-bridges Arg75-Asp155 and Arg88-Asp157 in the RRM1-RRM2 interface, which have been observed in all reported structures of free and nucleic-acid bound forms of UP1^24,23,25,26^. Introducing charge clashes (UP1-Arg75Glu/Arg88Glu) in this interface, which is remote from the RNA binding surface, decreases the binding affinity to the 12-mer RNA by ≈3-fold (*K*_D_ ~15.5 nM vs. ~40 nM) (Supplementary Fig. 2c). This suggests that the salt bridges play an indirect role for RNA binding by stabilizing the arrangement of the two RRM domains. To assess the effect of RNA-binding in the functional activity of UP1, we mutated conserved Phe residues within RNP-1 motifs that directly contact the RNA and are required for RNA-binding of hnRNP A1^21^. Substitution of Phe with Asp or Ala within individual or combined RRM1 and RRM2 domains was sufficient to abolish the activity of UP1-M9 in our *in vivo* pri-mir-18a processing assay without affecting the nuclear localization of the protein constructs (Supplementary Fig. 1a, b). This indicates that the RNA-binding activity of hnRNP A1 is essential for its stimulating activity of miRNA-18a biogenesis. The *in vivo* functional data further confirm that the stimulatory function of hnRNP A1 in processing of pri-mir-18a requires both RRM domains, as mutations that affect RNA binding in one domain (or deletion of one domain) abolish the activity of hnRNP A1.

Collectively, the structural model of the 1:1 UP1/12-mer RNA complex is fully consistent with our biochemical and functional data regarding the requirement for two RRM domains and two UAG motifs for high-affinity interaction.

### UP1 binding destabilizes the dynamic pri-mir-18a RNA terminal loop

The structure of UP1 bound to the 12-mer RNA in a 1:1 stoichiometry implies that the loop-proximal region of the pri-mir-18a stem should be destabilized to enable recognition of two UAG motifs, one accessible in the terminal loop and the second one as part of the pri-mir-18a duplex. To study this further, we first analyzed the structure of pri-mir-18a alone. For this, a model of the RNA was prepared using the MC-Fold/MC-Sym server^31^ (Fig. 4a) and assessed with experimental SAXS and NMR data. The predicted secondary structure and elongated shape of pri-mir-18a 71-mer is supported by SAXS data of the free RNA (Fig. 4b). We then used NMR to analyze the base-pairing in the pri-mir-18a 71-mer using imino NOESY spectra (Fig. 4c), as imino proton NMR signals probe the presence and stability of base pairs. We could unambiguously assign imino-imino cross-peaks corresponding to the stem region of the 71-mer RNA. However, no imino correlation was observed for the upper part of the stem loop (Fig. 4c, see secondary structure on the right), suggesting that this region of the RNA helix is dynamic with less stable base-pairing.

This was further confirmed by analyzing a 17-mer stem-loop construct (Fig. 4d, e), which represents the loop-proximal stem region in the full-length pri-mir-18a (Fig. 1b). The upper stem region in this RNA harbors the second UAG motif that is present in the single-stranded 12-mer RNA and recognized by UP1. An imino NOESY spectrum of the 17-mer RNA shows imino correlations only for the G:C and G:U base pairs indicating that the predicted A:U base pairs adjacent to the terminal loop are dynamic (Fig. 4d). Nevertheless, the presence of the A:U base pairs is confirmed by the (Py) H(CC)NN-COSY experiment, which can detect transient and weak-pairs^32^ (Fig. 4e). The absence of detectable imino protons for many predicted base-pairs in the 17-mer and in the upper part of pri-mir-18a indicates that the corresponding stem region exhibits weak base pairs and is dynamic.

The observation of dynamic base-pairing in the loop-proximal region is consistent with melting of the stem region to enable recognition of both UAG motifs by UP1. In fact, NMR titrations of UP1 with the 17-mer stem-loop RNA show virtually identical CSPs as seen with the 12-mer RNA (Fig. 2d, Supplementary Fig. 3c). As NMR chemical shifts are sensitive indicators of the three-dimensional structure, this finding confirms that the protein-RNA interactions observed with the 12-mer RNA also reflect the RNA recognition within the 17-mer RNA. In both cases, two UAG motifs are involved in the interaction, thus requiring melting of the 17-mer RNA helical stem. To further support this, we studied an A35C variant of the 17-mer RNA, which introduces complete Watson-Crick complementarity in the RNA stem. In contrast to the 17-mer, the 17-mer (A35C) exhibits all expected base-pairs as evidenced by detectable imino protons throughout the stem region (Fig. 4f). Consistently, NMR titrations show much smaller chemical shift perturbations in UP1 for the mutant 17-mer RNA (Supplementary Fig. 3d), which binds with a *K*_D_ of ~3 µM, i.e. corresponding to a 20-200-fold reduced binding affinity compared to the full-length pri-mir-18a and the single-stranded 12-mer RNA (Table 1; Supplementary Fig. 2d). Thus, the availability of only one single-stranded UAG motif for binding to UP1 yields micromolar affinity comparable to the 7-mer RNA. The fact that pri-mir-18a exhibits one UAG motif in the terminal loop and a second one in the weak and dynamic upper stem region suggests that binding of UP1 will require melting of the stem region flanking the terminal loop to enable recognition of this partially hidden UAG motif and high affinity (low nanomolar *K*_D_) RNA binding.

Next, we compared the ^1^H, ^15^N imino correlations of the pri-mir-18a 71-mer RNA in the free form and bound to UP1 (Fig. 4b). Notably, signals for the U10:G52 base pair in the middle of the stem region are not detectable in the complex but readily observed in the free form, indicative of destabilization and partial melting of this part of the duplex. Also, other residues especially next to mismatches and other less stable regions of the RNA exhibit reduced intensity or chemical shift perturbation such as G23, U49 and U59 (Fig 4c, green nucleotides). This is consistent with allosteric effects that lead to destabilization of the complete pri-mir-18a stem-loop induced by binding of UP1 to the terminal loop.

### Structural model of the UP1/pri-mir-18a complex and biochemical validation

To derive a structural model of UP1 bound to pri-mir-18a, we performed molecular dynamics simulations restrained by the structural information obtained for the UP1/12-mer RNA complex and experimental SAXS data of the UP1/pri-mir-18a complex (Fig. 5a; see Methods for details). The resulting structure was then scored against a combination of small angle neutron scattering (SANS) experiments with contrast matching (Fig. 5a; Supplementary Fig. 4e, f). SAXS data show that the dimensions of the complex correspond to a radius of gyration (R_g_) and maximum pairwise distance (D_max_) values of 37.7 Å and 130 Å, respectively, consistent with a 1:1 complex (Supplementary Table 2). The structural model shows very good agreement with all experimental data and is also consistent with *ab initio* SAXS derived models of the protein-RNA complex (Supplementary Fig. 4e, f). The UP1/pri-mir-18a complex shows the recognition of two UAG motifs in the terminal loop and a partially melted upper stem of pri-mir-18a (Fig. 5a).

**Fig. 5.**
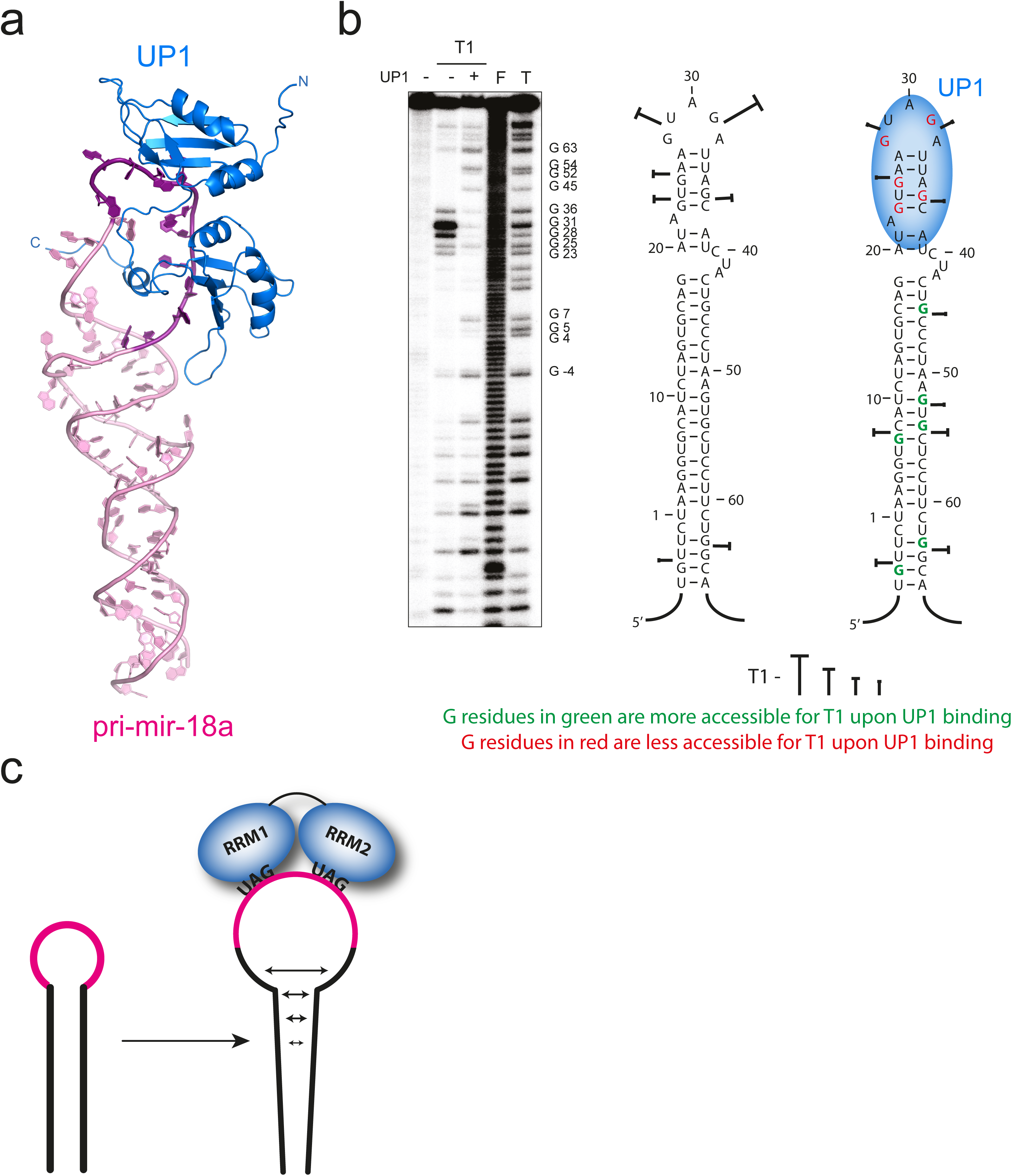
hnRNP A1 recognizes the terminal loop of pri-mir-18a. (**a**) Structural model of the 71-mer pri-mir-18a/UP1 complex. The region corresponding to the 12-mer RNA in the terminal loop of the pri-mir-18a is shown in dark magenta. (**b**) Footprint analysis of the pri-mir-18a/UP1 complex. Cleavage patterns were obtained for 5’ ^32^P-labeled pri-mir-18a transcript (100×10^3^ c.p.m.) incubated in the presence of recombinant UP1 protein (+ 200ng, 500nM), treated with Ribonuclease T1 at 1.5 U/μL. F and T identify nucleotide residues subjected to partial digest with formamide (every nucleotide) or ribonuclease T1 (G-specific cleavage), respectively. The cleavages intensities generated by Ribonuclease T1 are indicated on the pri-mir-18a secondary structure. The region of the major UP1 footprints is indicated by a blue oval shape. (c) A schematic model of the mechanism by which hnRNP A1 facilitates pri-mir-18a processing. Binding of UP1 to the terminal loop, where each RRM domain recognizes a UAG motif, leads to partial opening of the terminal loop that then spreads down into the RNA stem, thereby facilitating Drosha cleavage.

To validate the structural model described and to assess changes in accessibility of the pri-mir-18a induced by UP1 binding, we performed foot-printing analysis of the complete pri-mir-18a RNA in the absence and presence of UP1. This revealed that in the free RNA the terminal loop and flanking stem region comprising the two UAG motifs are accessible and dynamic (Fig. 5b), which is consistent with the NMR data (Fig. 4c, d). Binding of UP1 protects this region in the RNA, in full agreement with the structural model of the UP1/pri-mir-18a complex. The significant reduction in accessibility observed for residues in the terminal loop is consistent with the simultaneous binding of both RRM domains to the RNA (Figs. 3c, 5a). Interestingly, residues at the bottom part of the stem of pri-mir-18a become more accessible for nuclease cleavage upon binding of UP1 to the terminal loop (Fig. 5b, green nucleotides). This is consistent with the NMR analysis of pri-mir-18a free and bound to UP1 that shows UP1/pri-mir-18a interactions (Fig. 4) and the proposed destabilization of the RNA stem induced by UP1 binding (Fig. 5c).

### SHAPE analysis

Next, to assess the RNA structure of pri-mir-18a and potential effects from the presence of flanking regions in the context of the pri-mir-17-19 cluster, we performed structural analysis by selective 2’-hydroxyl acylation analyzed by primer extension (SHAPE, Supplementary Fig. 5a, b)^33^. To this end, *in vitro* transcribed RNAs comprising either pri-mir-18a or the pri-mir-17-19 cluster were incubated with increasing amounts of purified UP1 or full-length hnRNP A1 proteins, prior to treatment with N-methylisatoic anhydride (NMIA) that reacts with the 2´-hydroxyl group of flexible nucleotides. The SHAPE reactivity reflects the intensity of the NMIA-treated RNA primer extension products, normalized to the corresponding untreated RNA, in the presence or absence of UP1/hnRNP A1 proteins.

SHAPE analysis using pri-mir-18a indicates the presence of flexible regions in the terminal loop (nts 170-186) and one large bulge with increased accessibility (nts 148-149 and 211-218). Note that the numbers correspond to the pri-mir-17-19 transcript, with U152 in pri-mir-17-19 corresponding to U1 in pri-mir-18a (71-mer), whereas the 12-mer sequence from the pri-mir-18a (71-mer) terminal loop (A27-A38) corresponds to A178-A189 in pri-mir-17-19. Importantly, upon addition of UP1 and hnRNP A1, SHAPE differences relative to the free RNA indicate that residues preferentially protected from NMIA attack include the terminal loop and flanking stem region (nts 170, 185-186) (Supplementary Fig. 5a). The SHAPE reactivity values observed with pri-mir-17-19 free RNA were used to compute differences in SHAPE reactivity observed upon addition of proteins. Gross modifications of SHAPE reactivity in the presence of hnRNP A1 revealed the protection of nucleotides spanning the terminal loop of pri-mir-18a (nts 170, 177-186) and the region between miR-17 and miR-18a (nts 76, 94, 96, 102, 104-105, 113-116) (Supplementary Fig. 5b). Similar SHAPE results were observed upon incubation with UP1. Although the protection was less intense than that observed with the full-length protein, the protected residues mapped to the same RNA region (Supplementary Fig. 5). Importantly, these results confirm that the UP1 fragment of hnRNP A1 is sufficient to induce this protection, which is in agreement with the EMSA and functional assays (Fig. 1c, d). Most of the highly reactive residues in pri-mir-18a display a similar behavior as the pri-mir-17-19 transcript upon addition of UP1/A1 proteins (Supplementary Fig. 5). Despite the overall similar reactivity pattern, differences observed at nts 210-220 are presumably induced by the presence of the miR-17 and miR-19a in the whole transcript, which may stabilize the structure of the basal region of miR-18a. In summary, relative to the SHAPE reactivity observed with free RNA incubation of RNA with either hnRNP A1 or UP1 leads to a decreased SHAPE reactivity around the terminal loop of miR-18a, indicating that this region is the major binding site for the RRMs of hnRNP A1 and that this binding impacts the overall structure of pri-mir-18a in isolation or as part of the pri-mir-17-19 cluster.

### Mechanism of hnRNP A1 stimulation of pri-mir-18a processing

Finally, we attempted to address the mechanism by which hnRNP A1 activates the processing of pri-mir-18a. One possible scenario is that binding of hnRNP A1 (or UP1) leads to partial opening/melting of the terminal loop, which can lead to destabilization of the stem region and thus render it more accessible for processing by Drosha as proposed before^9^. Indeed, footprinting and site-directed mutagenesis of pri-mir-18a suggested that hnRNP A1 alters the local conformation of the stem in the vicinity of Drosha cleavage sites^9^, although the molecular mechanism was unclear. Indeed, UP1 (unwinding protein 1), as the name suggests, can unwind secondary and higher order structures of DNA and RNA^34,35,19^. To examine this possibility, we constructed a series of mutants in the terminal loop region of pri-mir-18a. These include single and double nucleotide mutants, where an A residue within the UAG motif in the terminal loop or within the second UAG motif, was mutated to C (UCG within pri-mir-18a[A30C] and pri-mir-18a[A30C/A35C], respectively), and a triple mutant (pri-mir-18a[U21A/U29A/U34A]), where all UAG motifs were mutated to AAG. We also designed a pri-mir-18a mutant, in which the terminal loop was stabilized by five G:C base pairs (pri-mir-18a[5GC]). As expected the wildtype sequence was efficiently processed. The single (pri-mir-18a [A30C]) and double nucleotide (pri-mir-18a[A30C/A35C]) terminal loop mutant RNAs retained hnRNP A1 binding, although with lower affinity, and were accordingly efficiently processed (Fig. 6a). The triple mutant (pri-mir-18a[U21A/U29A/U34A]) showed binding to hnRNP A1, although with reduced affinity, and retained efficient processing (Fig. 6b). The pri-mir-18a[5GC] mutant does not bind to hnRNP A1 in the RNA pull-down assay and consequently is not processed (Fig. 6a). This lack of processing could result from disruption of UP1 binding and/or conformational changes that inhibit Microprocessor activity, for example, by stabilizing the loop-proximal stem region. Based on these data we conclude that hnRNP A1 binding is essential for pri-mir-18a processing. The experiments also show that even very low hnRNP A1 levels are sufficient to stimulate processing activity.

**Fig. 6.**
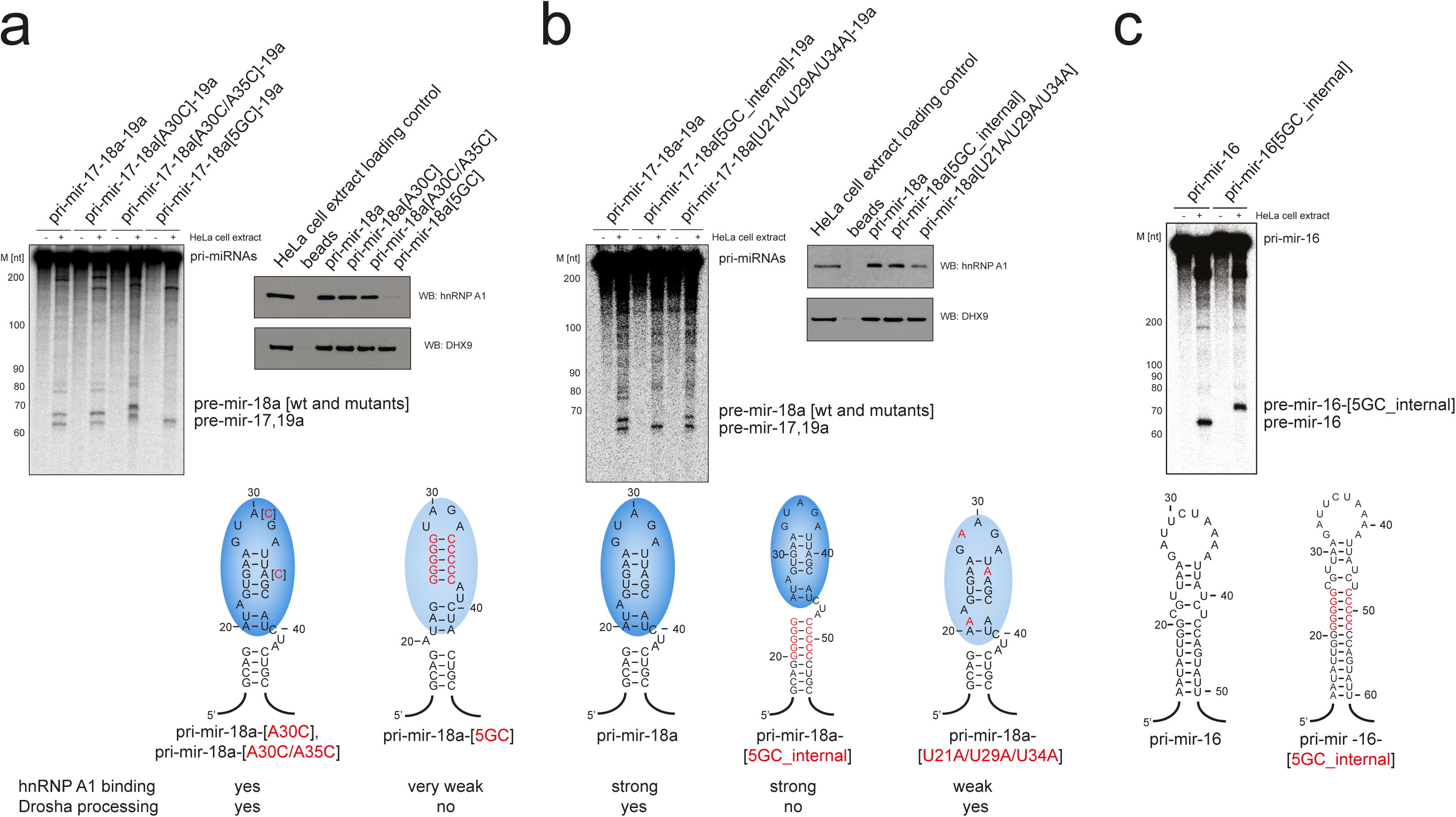
Mechanism of hnRNP A1 stimulation of pri-miRNA processing. (**a**) Pri-mir-18a with point mutations that were designed to weaken hnRNP A1 binding still binds hnRNP A1 in RNA pull-down assays and is processed by Drosha (*in vitro* processing assay with the pri-mir-17-19 cluster, wildtype and mutants), whereas pri-mir-18a mutant with a 5GC clamp does not bind hnRNP A1 (RNA pull-down assay is shown on the right) and is not processed by Drosha. (**b**) Pri-mir-18a with a 5GC_internal clamp and wildtype terminal loop binds hnRNP A1 but is not processed by Drosha. Pri-mir-18a with triple mutations [U21A/U29A/U34A] binds hnRNP A1 with lower affinity than the wildtype pri-mir-18a but is still efficiently processed by Drosha. (**c**) *In vitro* processing assay of pri-mir-16 with 5GC_internal clamp shows efficient processing by Drosha, similar to pri-mir-16 wildtype.

Next, we hypothesized that destabilization and unfolding of the upper stem region close to the terminal loop in pri-mir-18a RNA induced by hnRNP A1 binding can spread along the RNA stem and thus lead to destabilization of the RNA near the Drosha cleavage site. To examine this hypothesis we tested a different mutant pri-mir-18a, in which the terminal loop was clamped by 5 G:C base pairs (5GC_internal). Note, that this RNA variant is different from the pri-mir-18a[5GC] mutant described above, in which the terminal loop region was stabilized by 5 G:C base pairs (pri-mir-18a[5GC]). Remarkably, the processing of this RNA was impaired, despite retaining full binding to hnRNP A1 in the RNA pull-down assay (Fig. 6b). To rule out the possibility that the internal 5GC affects Drosha processing irrespective of the requirement for hnRNP A1 as an auxiliary factor, we examined the processing of pri-mir-16 with and without the 5GC clamp. Both wildtype pri-mir-16 and pri-mir-16[5GC_internal] were processed by Drosha, indicating that the internal 5GC clamp is not sufficient to impair Drosha processing (Fig. 6c). These data strongly support our hypothesis that unwinding of pri-mir-18a by hnRNP A1 can spread from the terminal loop towards the stem and is essential for stimulation of miR-18a biogenesis.

## Discussion

Here, we have used a multi-disciplinary approach to reveal the molecular mechanism by which hnRNP A1 binds to pri-mir-18a and facilitates its processing. Our results establish that hnRNP A1 specifically binds to pri-mir-18a through interactions involving both RRM domains in UP1 and a region comprising two UAG RNA sequence motifs in the terminal loop of pri-mir-18a and a flanking stem region. Cooperative binding of both domains to cognate RNA motifs results in substantially increased binding affinity and allows the unwinding of the target stem-loop RNA. This mode of binding is distinct from the recognition of a viral RNA, where apparently only RRM1 is involved in the recognition of a single AGU motif^25^, but reminiscent of recognition of two RNA motifs in single-stranded *cis* regulatory elements in alternative splicing regulation by hnRNP A1^26^. A common feature for all structures is that the overall domain arrangement and interface in free and nucleic acid bound hnRNP A1 tandem RRMs is conserved (Supplementary Fig. 4d). Nevertheless, some adaption and fine-tuning of the domain arrangement occurs upon RNA binding. This is also indicated by line-broadening of amide signals in the RRM1/RRM2 interface upon binding to the 12-mer RNA (Fig. 2e). A further common feature is the formation of 1:1 complexes in solution, very distinct from the 2:2 stoichiometry observed in crystal structures. The various distinct and specific modes of nucleic acid recognition by hnRNP A1 are intriguing and may reflect how it can play important roles in the regulation of many distinct biological activities by its RNA interactions.

hnRNP A1 can also act as a negative regulator of let-7a processing, by competing with the activator protein KSRP for binding to the pri-let-7a terminal loop^36,37^. In addition to hnRNP A1, several other RBPs recognize the terminal loop of miRNA precursors and influence either positively or negatively their biogenesis at the post-transcriptional level^38^. It was proposed that binding of Lin28 to the terminal loop of pre-let-7 leads to partial melting in the upper part of the stem, which can inhibit processing by both Drosha and Dicer^39^. Recently, Varani and coworkers showed that binding of the splicing factor Rbfox2 to the terminal loop of a subset of pri-miRNAs affects the conformation of the stem-loop structure and suppresses their nuclear processing^40^.

However, an important question is then how binding of a regulator to the terminal loop can affect Drosha cleavage at the opposite end of the RNA. Here, we propose that binding of hnRNP A1 to the terminal loop leads to destabilization of base pairs and (partial) melting of the loop-proximal stem region, with subsequent spreading of destabilization of the stem region towards the Drosha cleavage site (Fig. 5c). In support of this model, processing of a mutant pri-mir-18a, in which the terminal loop was clamped by 5 G:C base pairs (5GC_internal), was abolished, despite strong hnRNP A1 binding (Fig. 6b). This strongly argues that the effect of hnRNP A1 binding at the terminal loop is somehow propagated and leads to a stimulatory effect of Drosha processing.

We had previously shown that pri-mir-18b, which is part of the homologous primary cluster miR106a~18b~20b located on chromosome X, does not require hnRNP A1 for efficient processing. Mechanistically, this can be explained by the fact that the conformation of the stem in pri-mir-18b, resembles the more open stem structure comprising a bulge in the stem (UC→GU), which is only observed in pri-mir-18a, upon binding of hnRNP A1^8,9^. This is critical for more efficient Drosha processing as shown by the fact that simply introducing this bulge in the pri-miR-18a stem (UC→GU) made its processing more efficient and completely independent of the presence of hnRNP A1^8,9^.

Importantly, our use of integrative structural biology combined with biochemical and functional assays allowed us to extend these previous observations and conclude that the main effect of hnRNP A1 binding to the terminal loop of pri-mir-18a is to promote the destabilization of the lower stem, which leads to increased Drosha cleavage, via a mechanism that is not fully understood. Recent biochemical and structural analyses have shown that the Microprocessor recognizes two regions at either end of the miRNA precursor^41^. We found that strengthening the upper part of pri-mir-18a stem by GC base-pairs blocks miRNA processing, whereas disruption of the base-pairs enhances Microprocessor cleavage efficiency. It is noteworthy, that the partial unwinding of the apical RNA helix by binding of hnRNP A1 induces an asymmetry in the stem region that may define in which orientation the pri-mir-18a is recognized by the Microprocessor.

The processing efficiency of pri-mir-18a is context-dependent, suggesting that the sequence and/or structure of 18a as part of the miR-17-92 cluster is not optimal for Drosha processing^8^. Interestingly, several studies have recently shown that the miR-17-92 cluster adopts a compact tertiary structure, in which individual miRNAs have different expression levels depending on whether they are located on the surface or buried inside the core^42–44^. Notably, a recent SHAPE analysis of the miR-17-92 cluster revealed that the terminal loop of pri-mir-18a in the cluster is solvent inaccessible^42^. As the terminal loop corresponds to the sequence that we have identified as the main hnRNP A1-binding site in pri-mir-18a, it is tempting to speculate that binding of hnRNP A1 to the miR-17-92 cluster is associated with a conformational change in the RNA, which can facilitate Drosha cleavage. We propose that the tertiary structure of pri-mir-18a in the context of the miR-17-92 cluster, as well as sequences in the stem and loop region are important determinants of miRNA processing by Drosha. This process is regulated by the trans-acting factor hnRNP A1, which primarily interacts with the conserved terminal loop of pri-mir-18a. The recognition of the pri-mir-18a stem-loop by hnRNP A1 thus adds an additional layer for the regulation of pri-miRNA processing by an RBP, beyond features that have been recently identified^45–47^.

In conclusion, our data demonstrate that recognition of a conserved terminal loop RNA sequence in pri-miRNAs by an RBP can strongly modulate miRNA biogenesis by conformational changes and dynamic destabilization induced by RNA binding. Together with few recent reports, this suggests that recognition of pri-miRNAs by RBPs is a general paradigm for context-dependent regulation of miRNA biogenesis and function.

## Methods

### Protein expression and purification

The genes encoding RRM1 (1-97), RRM2 (94-196) and UP1 (1-196) were cloned into the pETM11 vector (EMBL) encoding a His tag followed by TEV cleavage site. Recombinant proteins were expressed in *E. coli* BL21(DE3) cells in standard media or minimal M9 media supplemented with 1 g/L ^15^N-ammonium chloride and 2 g/L ^13^C-glucose. Protein expression and purification was done as described previously^12^. After growth of bacterial cells up to an OD_600_ of 0.7-0.8, protein expression was induced by 0.5 mM IPTG and continued overnight at 20°C. Cells were resuspended in buffer A (30 mM Tris/HCl pH 7.5, 500 mM NaCl, 10 mM imidazole, 1 mM TCEP, 5% (v/v) glycerol) supplemented with protease inhibitors and lysed by sonication. The cleared lysate was loaded on Ni–NTA resin and after several washing steps with buffer A and buffer A containing 25 mM imidazole the protein was eluted with elution buffer containing 300 mM imidazole. After cleavage of the tag by His–tagged TEV protease at 4°C overnight, samples were reloaded on Ni–NTA resin to remove the tag, TEV protease and uncleaved protein. All protein samples were further purified by size-exclusion chromatography on a HiLoad 16/60 Superdex 75 column (GE Healthcare) equilibrated with NMR buffer (20 mM sodium phosphate pH 6.5, 100 mM NaCl, 2 mM DTT).

### Expression vectors

The plasmid pCGT7 hnRNP A1 and pCG T7 UP1 have been previously described^48^. In brief, these expression vectors are under the control of the CMV enhancer/promoter and include an N-terminal T7 epitope tag that corresponds to the first eleven residues of the bacteriophage T7 gene 10 capsid protein. The construct pCG T7 UP1-M9 harbors the M9 import/export sequence fused downstream of both RRM domains of UP1. The UP1-M9 mutants were synthesized by Invitrogen including the corresponding mutations and subcloned into pCG T7 (Supplementary Table 1). The mutated residues correspond to F148D/F150D (UP1-M9-FD2); F57D/F59D/F148D/F150D (UP1-M9-FD12); F17D/F57D/F59D/F108D/F148D/F150D (UP1-M9-FD6); F17A/F57A/F59A/F108A/F148A/F150A (UP1-M9-FA6)

### Indirect immunofluorescence

Cells were fixed and permeabilized for immunofluorescence assays at 24 hr after transfection. Fixation was with 4% p-formaldehyde in PBS for 15-30 min at room temperature, followed by incubation for 10 min in 0.2 % Triton X-100. The fixed cells were incubated for 1 hr at room temperature with 1:1000 anti-T7 monoclonal antibody (Novagen Inc.), washed with PBS, and incubated for 1 hr at room temperature with 1:200 fluorescein-conjugated goat anti-mouse IgG (Cappel Laboratories). The samples were observed on a Zeiss Axioskop microscope and the images were acquired with a Photometrics CH250 cooled CCD camera using Digital Scientific Smartcapture extensions in software from IP Lab Spectrum. The immunofluorescence figures show representative data, and each experiment was reproduced in multiple independent transfections.

### Analysis of miRNA levels in living cells

HeLa cells were grown in standard DMEM medium (Life Technologies) supplemented with 10% FBS (Life Technologies). Plasmids were transfected into HeLa cells using Lipofectamine 2000 reagent, as previously described^49^. Mouse monoclonal anti-T7 tag HRP conjugate (1:10000, 69048, RRID – AB10807495, Novagen) was used to detect T7-tagged proteins. Mouse monoclonal anti-tubulin (1:10000, T6199, RRID – AB_477583, Sigma-Aldrich) was used as loading control. miRNA qRT-PCR analysis was performed using the miScript qRT-PCR kit (Qiagen) on total RNA isolated with TRIzol reagent (Life Technologies), and each sample was run in duplicate. To assess the levels of the corresponding microRNAs, values were normalized to 5S RNA. For each measurement, three independent experiments were performed.

### RNA samples

Unlabeled 7-mer, 10-mer, 12-mer and 17-mer RNA oligonucleotides were purchased from IBA in double-desalted form (Göttingen, Germany). All other RNA samples were made by *in vitro* transcription and purified by denaturing PAGE. RNA samples were heated to 95°C for 3 min and snap-cooled on ice before use to promote proper folding.

### EMSA

Electrophoretic mobility shift assays (EMSA) were performed with internally labeled transcripts and proteins produced in *E. coli*. Gel-purified probes (50×10^3^ c.p.m. (counts per minute), ~20 pmol) were incubated in 15-μl reaction mixtures containing the indicated amounts of proteins in Roeder D buffer (100 mM KCl, 20% (v/v) glycerol, 0.2 mM EDTA, 100 mM Tris at pH=8.0, 0.5 mM DTT, 0.2 mM PMSF) supplemented with 0.5 mM ATP, 20 mM creatine phosphate, and 3.2 mM MgCl_2_. Reactions were incubated at 4°C for 1 h followed by electrophoresis on a 6% (w/v) non-denaturing gel. The signal was registered with radiographic film or was exposed to a phosphoimaging screen and scanned on a FLA-5100 scanner (Fujifilm).

### In vitro processing assays

Pri-miRNA substrates were obtained by *in vitro* transcription with [alpha-^32^P]-UTP. Gel-purified substrates (20×10^3^ c.p.m. (counts per minute), ~20 pmol) were incubated in 30 μl reaction mixtures containing 50% HeLa cell extract in Roeder D buffer, 0.5 mM ATP, 20 mM creatine phosphate, and 3.2 mM MgCl _2_. Then the reactions were incubated at 37°C for 30 min. The reactions were stopped with 2x Urea Dye, and followed by 8% (w/v) denaturing gel electrophoresis. The signal was registered with a radiographic film or by exposure to a phosphoimaging screen and scanning on a FLA-5100 scanner (Fujifilm).

### RNA pull-down

RNA pull-down was performed as previously described^49^. In summary, protein extracts from HeLa cells were incubated with *in vitro*-transcribed RNAs chemically coupled to agarose beads. The incubation was followed by three washes with buffer G (20 mM Tris pH 7.5, 135 mM NaCl, 1.5 mM MgCl_2_, 10% (v/v) glycerol, 1 mM EDTA, 1 mM DTT and 0.2 mM PMSF). After the final wash, the proteins associated with the beads were analyzed by SDS–PAGE, followed by western blotting. The following antibodies were used: rabbit polyclonal anti-DHX9 (1:1000, 17721-1-AP, RRID – AB_2092506, Protein-Tech); rabbit polyclonal anti-hnRNP A1 antibody (1:1000, PA5-19431, Invitrogen).

### Footprinting assays

The assays were performed as described earlier^50^. In brief, the substrates were synthesized by *in vitro* transcription and were 5’ labeled with PKA. A formamide ladder and ribonuclease T1 ladder were generated to assign the position. Ribonuclease T1 at 0.5 U/μL was added to reaction with 1 μL of RNA (50×10^3^ c.p.m.) and 7 μL of 1x structure buffer (12 mM Tris-HCl at pH=7.5, 48 mM NaCl, 1.2 mM MgCl_2_). Samples were unfolded at 90°C for 1 min and left at RT for 5 min to refold. The reactions were incubated at 37°C for 10 min. Reactions were run in the presence and absence of the recombinant UP1 protein. Reactions were resolved on 10% polyacrylamide gel. The signal was registered with a radiographic film or via exposure to a phosphoimaging screen and then scanned on a FLA-5100 scanner (Fujifilm).

### NMR spectroscopy

NMR experiments were recorded at 298 K on 900, 800 and 600 MHz Bruker Avance NMR spectrometers, all equipped with cryogenic triple resonance gradient probes. NMR spectra were processed by NMRPipe^51^ and analyzed using Sparky (T. D. Goddard and D. G. Kneller, SPARKY 3, University of California, San Francisco). All NMR samples were in NMR buffer with 10% D_2_O added as lock signal. Backbone resonance assignments of UP1 alone and in complex with RNA were obtained from a uniformly ^15^N,^13^C-labelled UP1 (with random fractional deuteration) in the absence and presence of saturating concentrations of RNA. Standard triple resonance experiments HNCA, HNCACB and CBCA(CO)NH**^52^** were recorded at 600 MHz.^*15*^N relaxation experiments were recorded on a 600-MHz spectrometer at 25°C. ^15^N T1 and T1ρ relaxation times were obtained from pseudo-3D HSQC-based experiments recorded in an interleaved fashion with 12 different relaxation delays (21.6, 86.4, 162, 248.4, 345.6, 432, 518.4, 669.6, 885.6, 1144.8, 1404 and 1782 ms) for T1 and 8 different relaxation delays (5, 7, 10, 15, 20, 25, 30 and 40 ms) for T1ρ. Two delays in each experiment were recorded in duplicates for error estimation. Relaxation rates were extracted by fitting the data to an exponential function using the relaxation module integrated in NMRViewJ^53^.

***Paramagnetic relaxation enhancements (PREs)*** were recorded at 600 MHz using a sample with concentration of ~200µM. UP1(E66C) was spin-labeled by adding 10-fold excess of 3-(2-Iodoacetamido)-PROXYL to the protein solution (in 100mM Tris pH 8.0, 100mM NaCl) and incubating at 4°C overnight. Unreacted spin-label was removed by size-exclusion chromatography. Completion of the spin-labeling reaction and attachment of a single spin-label on the protein was confirmed by LC/MS. PRE data were recorded and analyzed as described previously^28,54^. ^1^H^N^-^15^N ***residual dipolar couplings (RDCs)*** were measured on a ~200µM UP1/12-mer sample aligned in Pf1 phage (~10mg/mL) using the IPAP experiment^55^. RDCs were best-fitted to the structure by singular value decomposition (SVD) using PALES^56^.

### X-ray crystallography

Screening for crystallization conditions was done using commercial screens (QIAGEN and Hampton Research) at 20°C. UP1 (in 20 mM HEPES pH 7.0, 100 mM NaCl, and 1 mM TCEP) was mixed with 12-mer RNA in a 1:1 molar ratio (0.95 mM protein-RNA complex concentration) and incubated on ice for 1 h. Crystallization drops were set up by mixing 100 nL of complex and 100 nL of reservoir using the sitting drop vapor diffusion method. Crystals with hexagonal plate morphology were obtained in 0.2 M sodium citrate and 20% PEG 3350 after one week. The crystals were cryoprotected in reservoir solution supplemented with 20% (v/v) ethylene glycol and flash frozen in liquid nitrogen. X-ray diffraction data were collected at the European Synchrotron Radiation Facility (ESRF). All diffraction images were processed by XDS^57^. The CCP4 package^58^ was used for all subsequent data analysis. The structure of the UP1/12-mer RNA complex was solved by molecular replacement with the program Phaser^59^ using the coordinates of UP1 bound to modified telomeric DNA (PDB code: 1U1R)^60^ as search model. Model building was performed manually in Coot^61^ and refinement was done using Phenix^62^.

### Restrained molecular dynamics simulations

MD simulations were performed using GROMACS^63^ and PLUMED^64^. The systems were prepared using the AMBER99SB-ILDN force field in tandem with the TIP3P water model. After solvation in a cubic box, a short initial energy minimization of 100 steps was performed to resolve steric clashes between solvent atoms and the complex. Structures were refined using simulated annealing simulations. The temperature was adjusted with a period of 20 ps between 300 K and 100 K. The SAXS dataset of the UP1/12-mer RNA complex encompassed data points with scattering wavenumbers between 0.025 Å^−1^ and 0.685 Å^−1^. Intensities up to 0.5 Å^−1^ were fitted with a polynomial of 16^th^ degree. From this fit, 43 intensities were calculated for wavenumbers between 0.03 and 0.45 Å^−1^ in steps of 0.01 Å^−1^. These representative intensities were utilized as restraints. For the complex between full-length pri-mir-18a and UP1 representative intensities between 0.08 Å^−1^ and 0.22 Å^−1^ were calculated and used as restraints as described above. An approximate scaling factor relating calculated and measured SAXS intensities was estimated by the average ratio between the experimental SAXS intensities as taken from the fit and the first round of calculated SAXS intensities. SAXS intensities were calculated using the Debye formula and standard atomistic structure factors corrected for the effect of the solvent^65^.

Metainference^66^ with a Gaussian likelihood per data point on the representative SAXS intensities was applied every 10th step. It is of notice that metainference in the approximation of the absence of dynamics as used here (without replicas and a standard error of the mean set to zero) is equivalent to the Inferential Structure Determination approach^67^. The attributes of the uncertainty parameter were initially set to large values to allow a slow increase of the restrain force. Additionally, an additional scaling factor between the experimental data was sampled using a flat prior between 0.9 and 1.1.

The protein-RNA interface was restrained by harmonic upper-wall potentials centred 3.5 Å applied on the distances between the centres of the respective rings (Phe17 – A4, Phe59 – G5, Phe108 – A9 and Phe150 – G10) with force constants of 1000 kJ/mol. Furthermore, two crystallographic salt bridges (Arg75 – Asp155 and Arg88 – Asp157) between the two RRM domains were restrained with similar potentials centred at a distance of 4 Å applied on the distances between their charged groups. Secondary structures identified by STRIDE^68^ from the crystal structure were restrained using an upper wall potential on the rmsd of the backbone atoms of residues involved with a force constant of 10000 kJ/mol, centred at 0 Å.

For the UP1/12-mer complex RDCs restraints were applied using the θ-method^69^. To take into account for the multiple possible alignments of the molecule with the phage, RDCs were calculated as averages over two replicas and a linear restraint with a slope of −20000 kJ/mol was applied on the correlation between the average and experimental RDCs. Each replica was independently restrained with all the formerly introduced restraints.

After 300 preliminary annealing cycles, the refined structure was chosen as the one with the lowest metainference energy among those sampled at 300K in the latest 30 cycles. The quality of the structure was then further assessed using ProCheck^70^ and the SAXS/SANS profiles and RDCs were independently confirmed using Crysol^71^/Cryson^72^ and SVD, respectively.

### Isothermal titration calorimetry (ITC)

ITC measurements were carried out at 25°C using an iTC200 calorimeter (GE Healthcare). Both protein and ligand were exchanged into NMR buffer without DTT. Protein concentrations in the range of 300-1000 µM, depending on the affinity of the interaction, were injected into the sample cell containing RNA with a concentration of 20-100 µM. Titrations consisted of 20 injections of 2 µL or 26 injections of 1.5 µL with a 3-minute spacing between each injection. After correction for heat of dilution, data were fitted to a one-site binding model using the Microcal Origin 7.0 software. Each measurement was repeated at least three times.

### Static light scattering (SLS)

Static light scattering experiments were performed on a S75 10/300 size-exclusion column (GE Healthcare) connected to a Viscotek Tetra Detector Array (TDA) instrument equipped with refractive index (RI), light scattering, viscosity and photo diode array (PDA) detectors (Malvern Instruments). Sample volume of 100 µL was injected onto the column pre-equilibrated with NMR buffer and the flow rate was set to 0.5 mL/min. Calibration was done with BSA (bovine serum albumin) at a concentration of 4-5 mg/mL. Data were analyzed by the OmniSEC software using refractive index increment (dn/dc) values of 0.185 mL/g and 0.17 mL/g for protein and RNA samples, respectively^73^.

### Small angle X-ray/neutron scattering (SAXS/SANS)

SAXS data of UP1, pri-mir-18a and UP1/pri-mir-18a complex were recorded at the X33 beamline of the European Molecular Biology Laboratory (EMBL) at Deutsches Elektronen Synchrotron (DESY, Hamburg) at 15°C. The scattering curves were measured with 120-second exposure times (8 frames, 15 seconds each) for concentrations in the range of 1-10 mg/mL. The scattering intensity was measured covering the momentum transfer range 0.007 < q < 0.63 Å^−1^. Individual frames collected during the exposure time were compared to check for radiation damage before averaging. Scattering of buffer measured before and after each sample was averaged and subtracted from the scattering of the sample. SAXS data of the UP1/12-mer complex were recorded on a BioSAXS-1000 instrument (Rigaku) equipped with a Pilatus detector using 20 µL of sample in a capillary tube. All SAXS data were processed using the ATSAS software package^74^. The radius of gyration (R_g_) and the maximum dimension (D_max_) values were obtained from the GNOM program, which evaluates the pair-distance distribution function, P(r)^75^. For ab initio modeling of the protein-RNA complex, three scattering profiles corresponding to individual components and the complex were used as input for the multiphase modeling program MONSA^76^. Theoretical scattering curves were calculated using CRYSOL^71^.

SANS data were recorded at the large dynamic range diffractometer D22 at the Institut Laue-Langevin (ILL) Grenoble, France, using a neutron wavelength of 6 Å and a detector-collimator setup of 2m/2m. The scattering intensity was measured covering the momentum transfer range 0.02 < q < 0.35 Å^−1^. Sample and buffer volumes were 200 μL and exposed for 60 min. The 2D detector signals of the samples were corrected for detector efficiency, empty cell scattering, directly calibrated (against water), and azimuthally averaged using ILL in-house software (Gosh, R. E., Egelhaaf, S. U. & Rennie, A. R. *A Computing Guide for Small Angle Scattering Experiments*. Technical Report ILL06GH05T, 2006, Institut Laue-Langevin, Grenoble, France). The final 1D scattering curves were further analyzed using the ATSAS software.

### SHAPE analysis

Pri-mir17-19a (or pri-mir-18a) in complex with purified hnRNP A1 or UP1 proteins (125nM) were assembled in folding buffer (100 mM HEPES pH 8, 6 mM MgCl_2_) using 160 nM RNA prior to treatment with N-methylisatoic anhydride NMIA (Invitrogen), as the modifying agent, as recently described^77^. RNA was phenol extracted and ethanol-precipitated and then subjected to primer extension analysis. For primer extension, equal amounts of NMIA-treated and untreated RNAs (10 µL) were incubated with 0.5 µL of antisense 5′-end ^32^P-labeled primers 1-3, which were used for the analysis of regions comprising miR-17, miR-18a and miR-19a, respectively.

Primer1: 5´GCACTCAACATCAGCAGGCCCTGCAC 3’

Primer 2: 5´CTATATACTTGCTTGGCTTG 3´

Primer 3: 5´GACCTGCAGGCGGCCGCG 3´

Primer extension was conducted in a final volume of 15 µL containing reverse transcriptase (RT) buffer (50 mM Tris–HCl, pH 8.3, 3 mM MgCl_2_, 75 mM KCl, 8 mM DTT) and 1 mM of each dNTP. The mix was heated at 52°C for 1 min, prior to addition of 100 U of Superscript III RT (Invitrogen) and incubation at 52°C for 30 min. cDNA products were fractionated in 6% acrylamide, 7 M urea gels, in parallel to a sequence obtained with the same primer. For SHAPE data processing, the intensities of RT-stops were quantified as described^78.^. Data from three independent assays were used to calculate the mean (±SD) SHAPE reactivity.

**Supplementary Information** is linked to the online version of the paper.

## Acknowledgements

We are thankful to Marianne Keith (MRC HGU, University of Edinburgh) for Immunofluorescence data. We thank Alisha Jones for discussions. We are grateful to Robert Janowski for collecting the X-ray diffraction data and Ralf Stehle for help with in-house SAXS measurements. We thank the beamline staff at DESY (Hamburg) and Anne Martel at ILL (Grenoble) for support with SAXS and SANS measurements, respectively. We acknowledge the use of the X-ray crystallography platform at the Helmholtz Zentrum München, NMR access at the Bavarian NMR Center and SAXS measurements at the facility of the SFB1035 at Department Chemie, Technical University of Munich. This work was supported by the Deutsche Forschungsgemeinschaft through grants SFB1035 and GRK1721 (to M.S.), MRC core funding and the Wellcome Trust (Grant 095518/Z/11/Z to J.F.C.), Medical Research Council Career Development Award (G10000564 to G.M.). Wellcome Trust Centre Core Grants (077707 and 092076 to G.M.). M.M., A.J. and C.C. acknowledge the support of the Technische Universität München – Institute for Advanced Study, funded by the German Excellence Initiative and the European Union Seventh Framework Programme under grant agreement n° 291763.

## Author Contributions

HK performed molecular biology, protein/RNA preparation, ITC, NMR, crystallographic structure determination, RNA modelling and SAXS analysis; NRC and GM performed footprinting assays, NF performed SHAPE analysis, BS contributed to structural modeling, MM, AJ and CC performed structural modeling and restrained molecular dynamics, FG analyzed SAXS and SANS data, AD performed NMR of RNA samples. HK, GM, JFC and MS conceived and designed the project. HK, GM, JFC and MS wrote the paper. All authors discussed the results and commented on the manuscript.

## Author Information

Atomic coordinates and structure files for the UP1/12-mer RNA crystal structure have been deposited in the Protein Data Bank (http://www.pdb.org/) with accession code XXX.

The authors declare no competing financial interests.

